# Exploring the Knowledge of An Outstanding Protein to Protein Interaction Transformer

**DOI:** 10.1101/2023.02.09.527848

**Authors:** Sen Yang, Dawei Feng, Peng Cheng, Yang Liu, Shengqi Wang

## Abstract

Protein-to-protein interaction (PPI) prediction aims to predict whether two given proteins interact or not. Compared with traditional experimental methods of high cost and low efficiency, the current deep learning based approach makes it possible to discover massive potential PPIs from large-scale databases. However, deep PPI prediction models perform poorly on unseen species, as their proteins are not in the training set. Targetting on this issue, the paper first proposes PPITrans, a Transformer based PPI prediction model that exploits a language model pre-trained on proteins to conduct binary PPI prediction. To validate the effectiveness on unseen species, PPITrans is trained with Human PPIs and tested on PPIs of other species. Experimental results show that PPITrans significantly outperforms the previous state-of-the-art on various metrics, especially on PPIs of unseen species. For example, the AUPR improves 0.339 absolutely on Fly PPIs. Aiming to explore the knowledge learned by PPITrans from PPI data, this paper also designs a series of probes belonging to three categories. Their results reveal several interesting findings, like that although PPITrans cannot capture the spatial structure of proteins, it can obtain knowledge of PPI type and binding affinity, learning more than binary PPI.

## 1 Introduction

As the most common macromolecules in cells, proteins participate in most life activities through interacting with others (Protein-to-Protein Interaction, PPI). PPI is critical for various biological procedures, such as signal transduction [1] and regularization of metabolism [2]. Predicting PPIs could provide valuable perspectives for understanding cellular life activities.

Many traditional methods in the lab can detect PPIs, such as two-yest hybrid system [3] and protein chips [4]. However, all experimental methods to date share the disadvantages of labour consumption and low efficiency. So they can detect only a small fraction of true PPIs, being insufficient to handle the large-scale protein data generated by high-throughput sequencing. In contrast, computational methods for PPI prediction can overcome the disadvantages of experimental ones. They typically extract statistical features from existing PPIs and develop novel algorithms to discover numbers of unseen PPIs in the large-scale protein data at a low cost. So far, with the blooming of deep learning [5], numbers of deep models [6] have been proposed to solve the problem of PPI prediction. Since PPI prediction is comparable to sentence pair classification tasks in the natural language processing (NLP) [7], the deep models boosted PPI prediction by drawing successful cases in NLP, such as recurrent convolutional neural network [8] and siamese network [9].

Although the deep learning approach has made impressive progress in PPI prediction, there are still unsolved issues. One is that the deep models typically perform worse on unseen species. The proteins across species differ in residue sequence, structure and other properties. Figure 1 exposes such a gap by comparing protein identities of five species with the Human. It can be seen that most Mouse proteins show higher identities, indicating their homologous proteins can be found in provided human proteins. By comparison, protein identities of the other four species are generally low, especially E.coli, indicating that greater genetic distance can result in a larger difference gap. On the other hand, current deep PPI prediction models are typically trained on a species-specific PPI dataset that covers only a small fraction of existing proteins. Therefore, they are incapable of making trustworthy predictions in other species with a large difference gap of proteins [10]. This issue is also called the out of distribution (OOD) problem generally in the machine learning community. Solving the OOD problem could enable PPI prediction models to transfer effectively to other species and discover interaction pairs of unknown or designed proteins. Another issue is the interpretability of deep PPI prediction models. Recent works only focus on improving PPI prediction with deep models, giving little attention to their inside mechanisms, such as whether the model works as expected, what knowledge the model has learned and not learned. Answering these questions can promote our understanding to the underlying principles and reliability of deep PPI prediction models.

**Fig. 1.**
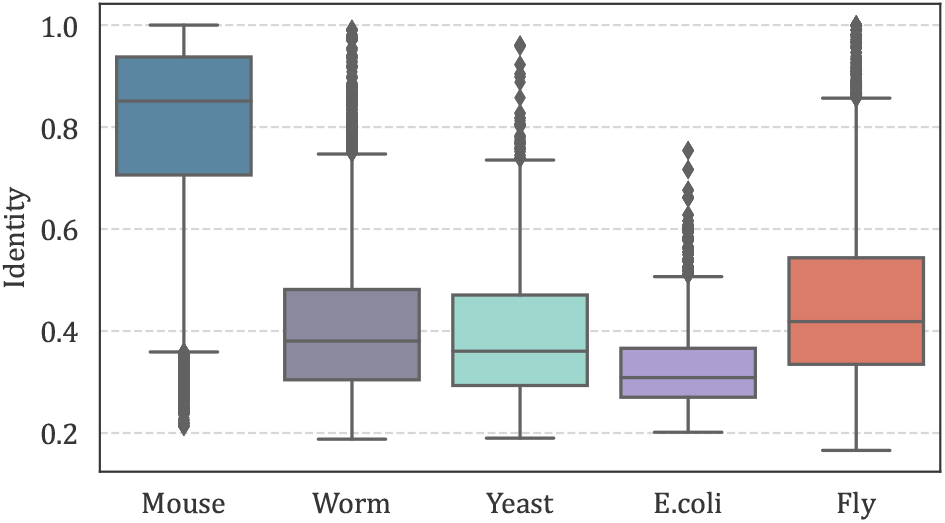
Maximum protein identites (computed by BLASTP) of five species with more than 70k Human proteins. The protein data is from https://github.com/samsledje/D-SCRIPT/blob/main/data/seqs.

Since the primary amino acid residue sequence has contained all necessary information for PPI [11] and remains available for almost all proteins, we focus on PPI prediction with only the primary residue sequences as in previous works [10], [12], [13], [14]. Targeting on the OOD problem, we first propose a Transfomer based PPI prediction model, namely **PPITrans**. PPITrans exploits a huge language model ProtT5 [15] as the feature extractor, which is a big language model pre-trained on proteins and surpasses other peer models on downstream tasks. Due to pre-trained on large-scale protein datasets [16] that cover many species, such a large model has valuable benefits in alleviating the OOD problem. On top of ProtT5, we adopt the siamese [9] architecture that contains two Transformers [17] for PPI prediction. We compare PPITrans with several strong baselines on the dataset [10] built from the STRING database [18]. Experimental results show that PPITrans significantly surpasses previous models on Human PPIs and has more evident advantages on PPIs of other species, although they are invisible to the model during training. For example, the AUPR of PPITrans improves 33.9 percentage points over the previous state-of-the-art on Fly PPIs.

Moreover, aiming to investigate the knowledge learned by PPITrans, we design a series of novel probes. Each probe is a task that aims to explore a specific knowledge of PPITrans. This probes can be classified into three categories according to the granularity. Overall, these probes are used to answer an interesting question not mentioned in previous works: *“What knowledge has the model learned and not learned in addition to the binary PPI?”*. Experimental results of these probes reveal several interesting findings: PPITrans gradually drop fine-grained residue features from the bottom layer to the top layer; Besides, although performing pretty well on PPI prediction, it seems that PPITrans makes predictions based on the co-occurrence of amino acids, as it cannot capture the spatial structure of protein or protein pairs; However, PPITrans can automatically learn more about protein pairs beyond binary interaction relations, such as PPI type and binding affinity, although they are not in the PPI dataset.

## 2 Related Works

This section presents recent efforts that have been made toward the development of deep learning methods for PPI prediction. As for other experimental or computational methods for PPI prediction, previous works have given detailed surveys [6], [19]. Existing deep learning methods for PPI prediction can be divided into three categories according to biological information fed to the model.

Because the amino acid sequences are sufficient for determining the interaction of two proteins [11] and they remain available in many databases [18], [20], [21], [22], most deep learning methods only rely on sequences to predict PPIs. DPPI [12] is an earlier model that employs a deep learning approach for binary PPI prediction. It applies a deep, siamese [9] convolutional neural network (CNN) to capture features from protein profiles. Although DPPI surpasses previous computational methods, it requires excessive effort to construct the protein profile, a matrix representing the probability that each amino acid appears in a specific position. DNN-PPI [14] uses CNN stacked with a long-short-term memory network [23] to extract protein feature. Unlike DPPI [12], the embeddings of amino acids are randomly initialized and optimized with the word2vec algorithm [24] in DNN-PPI [14]. PIPR [13] concatenates the pre-trained word2vec embedding and the one-hot encoding defined in [25] as the initial representation of amino acid. PIPR trains a recurrent convolutional neural network [8] of five layers with multiple learning objectives. However, stacking RNN of more than three layers will degrade the prediction performance as a result of gradient vanishing and exploding, which is already a consensus in the NLP community. D-SCRIPT [10], [26], like the above methods, also learns PPI prediction using only amino sequences. It adds a regularization term for every amino acid pair to learn the contact maps of protein pairs, which is computation inefficient.

The tertiary structure is also closely related to PPI [27]. As the AlphaFold [28] makes tertiary structures of more and more proteins available, several works have paid efforts to incorporate high-level structure information into a deep learning approach for PPI prediction [27], [29]. S2F [29] is a PPI prediction model pre-trained with sequences, structures, and functions of proteins. It takes the spatial structure as point clouds to model backbones and side chains. But the performance improvement on the PPI prediction task of S2F by pre-training is not significant. TAGPPI [27] integrates protein structures with amino acid sequences to improve PPI prediction. TAGPPI transforms the three-dimensional structure into a contact map matrix by setting a threshold of Euclidean distance and exploits the graph attention network [30] to encode the spatial structure. Although TAGPPI outperforms previous works, it lacks in-depth interpretation and analysis towards its learned knowledge.

PPIs can be combined to form a PPI network where nodes are proteins and edges denote interactions. The patterns of the PPI network can facilitate PPI prediction. For example, [31] argues that two proteins will interact if they have similar distances to other proteins in the PPI network. While [32] hypothesize that proteins interact if one of them is similar to the other’s partners. HO-VGAE [33] aggregates information of high-order neighbors into the representation of proteins in the PPI network. However, HO-VGAE cannot cope with query proteins that do not appear in the PPI network. GNN-PPI [34] integrates the sequence and partners into encoding for each protein to predict PPI of multiple types. This model takes all input protein pairs as potential PPIs so that it requires no negative samples for training. Such a setting makes GNN-PPI impractical to find true PPIs from not yet verified pairs.

Since the amino acid sequences of proteins determine their structures and functions, and they are available for almost all proteins, some recent methods are developed to intorduce pre-training, which have prevailed in the NLP field, into protein to predict biological functions. The earlier ones make use of word2vec algorithm [24] to embed protein sequences [35], [36]. However, their pre-trained models are either too shallow or trained with only a tiny fraction of existing proteins, being insufficient to boost model performance significantly. After that, large models are adopted. Unirep [37] explores the pre-training and finetuning paradigm. The following ESM-1b [38] also exploits such paradigm but is trained on a larger dataset. Prot-Trans [16] pre-trained many language models commonly used in the NLP field and compared their performance comprehensively. Since these deep models are pre-trained on large-scale protein sequences, they could capture the hidden patterns of natural proteins and benefit a lot of downstream tasks, such as protein function classification, secondary structure prediction, and protein sub-cellular location. Among these pre-trained models, ProtT5, one of ProtTrans [16] has been proven superior to others, including ESM-1b. Therefore, we build our model based on ProtT5 to promote PPI prediction.

## 3 Materials and Methods

### 3.1 Preliminary

We denote the training PPI dataset as 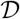. The *i*-th instance in 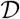 is represented as 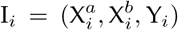, where *X* denotes protein, and *Y_i_* ∈ {0, 1} is a binary label indicating whether the two proteins interact (*Y_i_* = 1) or not (*Y_i_* = 0). We profile each protein as a sequence of amino acid residues. For example, 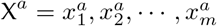. Here, 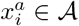 represents a residue that belongs to 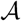, the set of 20 canonical amino acids. The PPI prediction model aims to learn a function *f* that maps a protein pair (*X^a^, X^b^*) to a binary label Y: *f*: *X^a^, X^b^* → Y. For a protein *X* = *x*_1_, *x*_2_,…, *x_m_*, the PPI prediction model firstly embeds it into vector representation 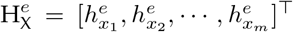, in which superscript *e* represents the embedding layer, H is the protein representation, and *h* is the residue vector. Since the embedding size differs from the hidden size of the protein encoder, there is a projection layer between them as described in Section 3.3.2. We denote the output of the projection layer as 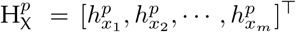, with superscript *p* representing the projection layer. We represent the output of *l*-th (1 ≤ *l* ≤ *L*) layer of the encoder as 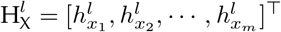.

### 3.2 Dataset

We use the PPI data of six species collected by [26] from STRING [18] to train and test PPI prediction models. The PPIs in the dataset are of various types, such as instantaneous interaction and permanently complex, while we leave out the PPI type since the PPI prediction is a binary classification task. Since our goal is to train a PPI prediction model for unseen species, PPIs of some species should be blind to the model during training. On the other hand, humans have the most annotated PPIs among all species. Therefore, to conduct cross-species PPI prediction, we take part of Human PPIs as the training set while the remaining PPIs as the test set. Actually, PPIs of other single or even multiple species can also serve as the training set, and the more training data the better in principle, However, splitting more species into the training set will reduce test results, thus not representative. In this split, there are about 27 thousand positive samples in the test set and 38 thousand in the train set. Except that E.coli has only 2000 positive samples, the positive samples for the test of other species are about 5000. Negative samples are generated following [12] by randomly pairing proteins from the non-redundant set. The number of negative samples is ten times that of the positive samples to reflect the intuition that true PPIs are rare. Table 1 shows details about the dataset.

**TABLE 1.** Statistics of the dataset.

| Species | Split | Positive | Negative | Proteins | Residues |
| --- | --- | --- | --- | --- | --- |
| Mouse | test | 5,000 | 50,000 | 40,606 | 15,933,279 |
| Fly | test | 5,000 | 50,000 | 19,310 | 7,491,553 |
| Worm | test | 5,000 | 50,000 | 25,930 | 9,121,823 |
| Yeast | test | 5,000 | 50,000 | 5,664 | 1,934,430 |
| E.coli | test | 2,000 | 20,000 | 17,722 | 5,076,871 |
| Human | test | 4,794 | 47,931 | 70,529 | 29,306,476 |
|  | train | 38,344 | 383,448 |  |  |

### 3.3 PPITrans

Figure 2 shows the architecture of PPITrans. It contains three modules: pre-trained embedder, protein encoder, and PPI classifier. The embedder exploits the pre-trained language model ProtT5 [16] to embed all input amino acid residues into vector space with position and context awareness. The protein encoder is a Transformer [17] that encodes embeddings output by the pre-trained embedder for the following PPI prediction. After encoding, a pooling layer turns vector representations of amino acids into a protein vector representation. Finally, the PPI classifier, a multilayer perceptron (MLP), takes the Hadamard product of two vector representations as input and outputs the interaction probability of the input protein pair.

**Fig. 2.**
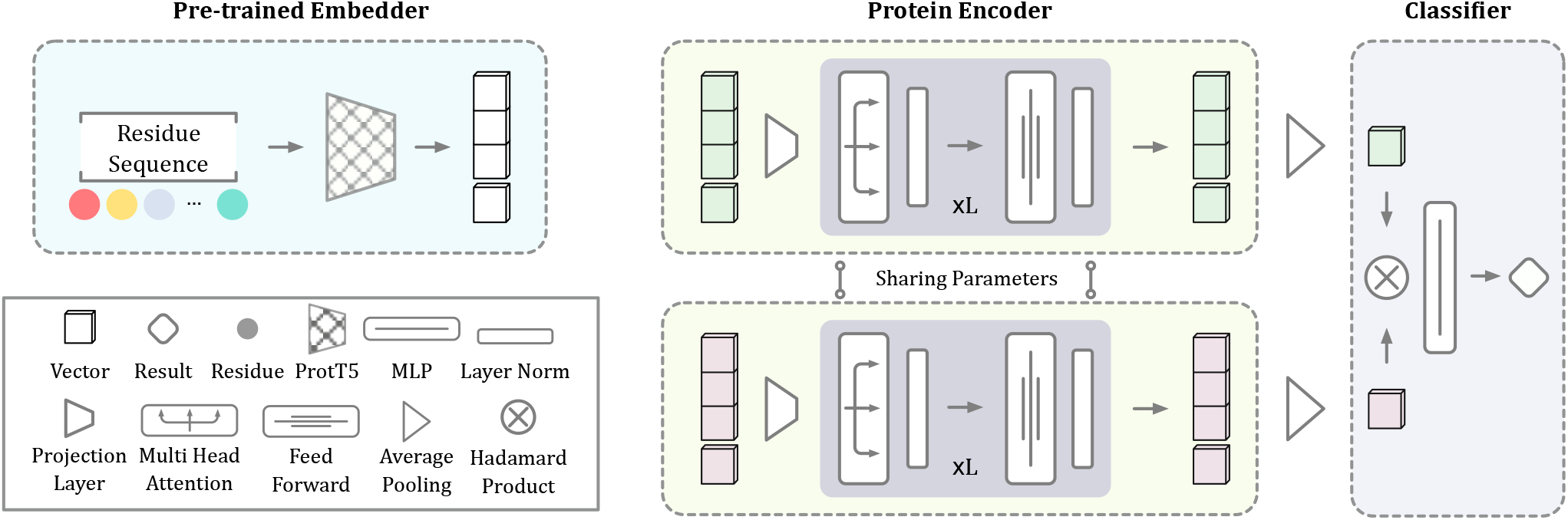
Overview of PPITrans. The lower left part is the legend, showing the representation of each symbol.

#### 3.3.1 Pre-trained Embedder

As pointed out above, one of our goals is to improve PPI prediction on unseen species. When a deep model trained on PPIs of a specific species transfers to another species of which the PPIs were not visible, its performance could degrade sharply as the large genetic distance between two species results in few similarities between their proteins.

Starting in the NLP field [39], pre-training has been proven beneficial to many research tasks. Some works tried to introduce pre-training into PPI prediction [26], [27]. However, their pre-trained models are either too shallow or trained with only a tiny fraction of existing proteins, being insufficient to boost model performance significantly [15]. Therefore, we try to alleviate the OOD problem by introducing a huge language model pre-trained on large-scale protein data to PPI prediction, which has not been investigated in previous works.

Specifically, we adopt ProtT5 [16] to embed amino acid sequences dynamically. The model containing 3 billion parameters is pre-trained on the BFD [40] dataset and finetuned on the UniRef50 [41] dataset which contains 45 and 2122 million proteins, respectively. After embedding, the *i*-th residue *x_i_* in a protein is embedded into a vector 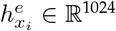.

#### 3.3.2 Protein Encoder

As ProtT5 requires excessive computation resources for finetuning, we freeze its parameters and stack a Transformer encoder on it to encode protein sequences further. Actually, many models can act as the sequence encoder, such as Transformer [17], RNN [23], and CNN [8]. Among these models, we select Transformer as the encoder for two considerations. Firstly, deep models typically have superior performance than those shallow. Then RNN based encoders are excluded because they are difficult to stack into the deep. Secondly, we aim to probe the knowledge of the model learned from three levels, one of which is the residue level. So the encoder should keep the sequence length of hidden outputs unchanged. Then CNN based encoders are excluded due to their disadvantage of changing sequence length. So we finally exploits the Transformer as the encoder.

The vector size reaches 1024 in the pre-trained embedder, being unsuitable for the encoder as it requires more computation resources without performance improvement. Therefore, we add a projection layer to transform the embedding size into the hidden size *d* of the Transformer encoder:

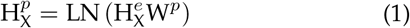

where 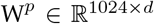 is a parameter matrix, and LN is the layer normalization function.

The Transformer encoder contains two conceptual modules inside each layer: the multi-head attention and feedforward network. The multi-head attention module comprises several independent self-attention modules to learn multi-view context aware representation, following a layer normalization and residual connection. It can be formulated as:

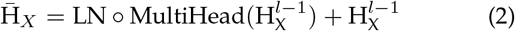

where 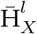 represents the output of multi-head attention module in the *l*-th layer, and ◦ denoes fuction composition. Note that 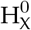 equals to 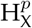 in the Equation 1. The MultiHead function in Equation 2 works as:

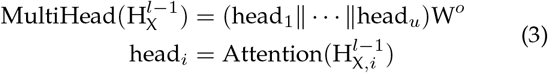

It concatenates outputs of all heads and multiplies with a parameter matrix 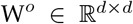. In multi-head attention, protein representation 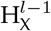 is sliced longitudinally into *u* heads. So 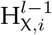 is the *i*-th splice of 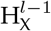. If 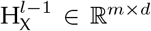 then 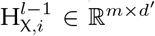 where 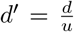. The Attention function in Equation 3 is the self-attention mechanism:

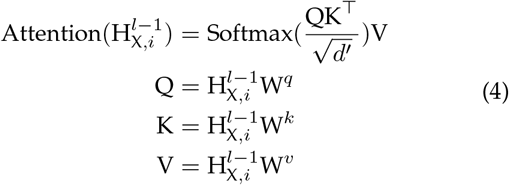

Here, 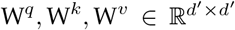 are three parameter matrices. Unlike the multi-head attention module that captures context information through self-attention operation, the feedforward module conducts extraction for each amino acid separately and identically. It contains two linear layers with Relu activation and also exploits layer normalization and residual connection:

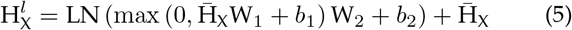

in which 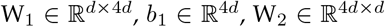, and 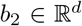 are all learnable parameters. 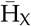 is the output of multi-head attention module computed as in Equation 2.

Since each input PPI sample contains two proteins, we adopt the siamese architecture [9] for protein encoding. The two encoders share parameters in the siamese architecture, as shown in Figure 2.

#### 3.3.3 Prediction Module

The classifier makes predictions according to the output of the protein encoder. Assuming the input two proteins are *X^a^* and *X^b^* of length *m* and *n*. Their representations outputed by the encoder are 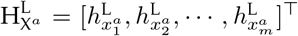 and 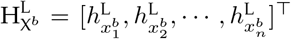. The prediction module first squeezes 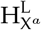 and 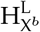 into two vectors through average pooling:

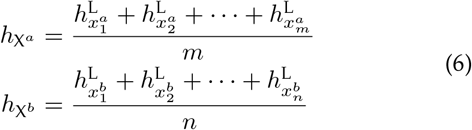

Then the the classifier outputs the probability distribution with a MLP:

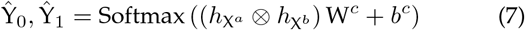

In Equation 7, ⊗ denotes the Hadamard product, 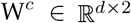 and 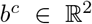 are parameters of the classifier module, while 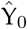 and 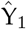 represent the probabilities of noninteraction and interaction, respectively. Through Equation 6&7, we can measure quantitatively the contribution of any two residue pairs to 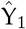. Specifically, 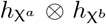 in Equation 7 is equal to:

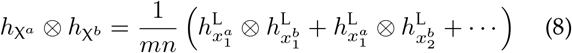

We then define two variables for any residue pair 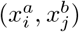 as:

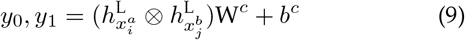

Taking Equation 8&9 into Equation 7, we can find that large *y*_1_ – *y*_0_ will result in large 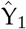, thus we define *y*_1_ – *y*_0_ as the contribution of a residue pair to the protein interaction.

In the traning stage, the objective of PPITrans is to minimize the cross-entropy between predcited probability and the gold label:

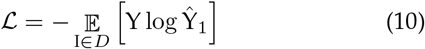

### 3.4 Probes

In this section, we devise a series of novel probes to explore the knowledge learned by PPITrans. As shown in Figure 3, each probe is a task that exploits classifiers or regressors to validate whether the representations output by each layer of PPITrans could provide specific information. Table 2 list all probes. According to the granularity, these probes can be classified into three levels: residue level, protein level, and protein pair level. Because PPITrans has ten layers (embedder layer, projection layer, and eight encoder layers), each probe has ten identical classifiers/regressors that work simultaneously. For probes of type classification, we use the K-Nearest Neighbors (KNN) with the nearest five neighbors as the classifier. As for the probes of regression type, we use the Kernel Ridge Regressor (KRR) to estimate the target variable. If we had used a more sophisticated classifier or regressor such as random forest and neural network, it would be unlikely to tell whether a certain result was due to the power of the PPITrans or some non-linear transformation performed by the classifier or regressor.

**Fig. 3.**
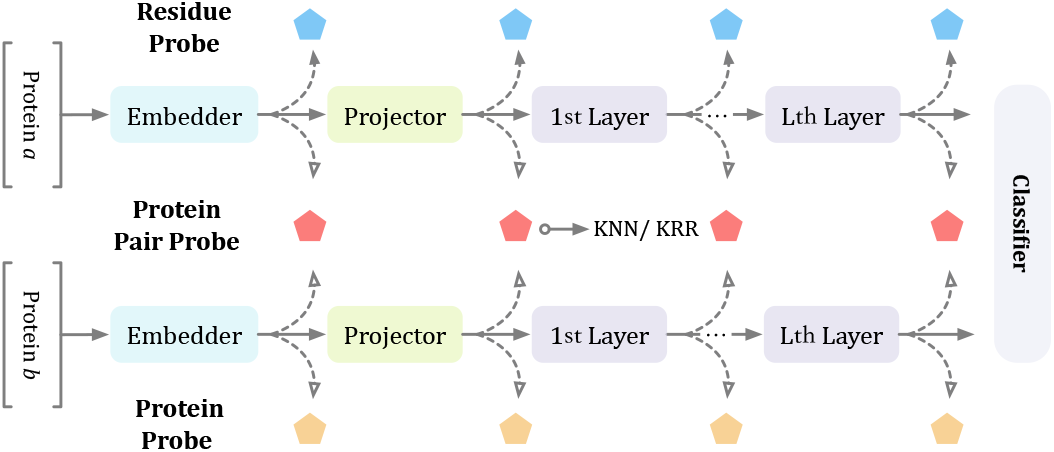
Overview of probes at three levels. The hollow arrow indicates that residue embeddings are transformed into protein embedding by average pooling.

**TABLE 2.** List of probes.

| Leve | Probe | Type |
| --- | --- | --- |
| Residue | Amino Acid Classification | Classification |
|  | Physicochemical Property Prediction | Classification |
| Protein | Secondary Structure Prediction | Classification |
|  | Contact Map Prediction | Classification |
|  | Species Classification | Classification |
| Protein Pair | PPI Type Prediction | Classification |
|  | Protein Pair Contact Map Prediction | Classification |
|  | Binding Affinity Estimation | Regression |

At the residue level, there are two probes: amino acid classification and physicochemical property prediction. They are used to identify the amino acid and the physicochemical property given the output of each layer. For the first one, we use a KNN to identify the amino acid from its representation. Specifically, the training and test set contains 168,521 and 43,071 (5 folds) amino acids, randomly selected from proteins in STRING [10], for the KNN classifier. As for the second probe, [25] divided 20 amino acids into 7 classes according to their physicochemical properties, so we use these 7 classes as the label of amino acids. This probe reuses the settings and data of the first one.

At the protein level, three probes aim to verify whether the representation contains various information about a protein, such as the secondary structure, species it belongs to, and contact map. The first probe targets on the secondary structure. We put this probe at the protein level because the secondary structure is determined by the whole residue sequence rather than just one residue. The protein secondary structure can also be represented as a sequence of characters, each indicating the local folded conformation of the corresponding amino acid. We randomly select 15,806 and 4,149 annotated residues as the training and test set from DSSP [42]. The probe of contact map prediction is to check whether the representation contains spatial information. Specifically, we use the contact map to represent the protein spatial structure and exploit KNNs to predict whether or not two residues in a protein are in contact. The contact map of a protein is determined as the following:

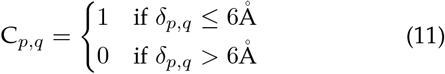

where C is the contact map, *δ* denotes Euclidean distance. Equation 11 means that two residues are in contact if the Euclidean distance between their alpha carbon atoms is less than or equal to 6Å. In this way, from AlphaFold database [28] we collect 112,935 residue pairs (22,587 in contact and 90,348 not in contact) as the training set and 23,110 residue pairs (4,538 in contact and 18,752 not in contact) as the test set. The last probe at the protein level aims to detect whether protein representations keep separated concerning species. We randomly select 12,000 proteins (2,000 for each species) from the dataset and split 9,600 for training and 2,400 for testing. As in previous probes, each layer is assigned a KNN to predict the species of given protein representations, which is the average pooling of its amino acid representations (Equation 6).

At the protein pair level, there are also three probes. They go further than binary PPI prediction and aim to explore fine-grained interaction details, including PPI type, contact map of protein pairs, and binding affinity. The first probe is to predict the contact relation between two residues of protein pairs. We construct data in a similar procedure as the probe of contact map prediction at the protein level (Equation 11), except that the threshold is set to 10Å. Specifically, we collect 155,510 residue pairs (31,102 in contact and 124,408 not in contact) as the training set and 26,860 residue pairs (5,372 in contact and 21,488 not in contact) as the test set from from GWIDD [43], which containing the docking information of interacted proteins. The second probe is to predict the type of interacted protein pairs. We use the SHS27k dataset [13] that contains 14,367 training and 4,789 testing PPIs of seven types. The KNN classifiers in this probe take the Hadamard product of two protein vectors (Equation 6) and output their interaction type. The last probe is to estimate the binding affinity of two interacted proteins. We use the SKEMPI dataset [13] that contains 2,213 training and 848 testing protein pairs with their binding affinities. In this probe, we assign a KRR to each layer and train them with the goal of minimizing the mean square loss (MSE).

## 4 Results

### 4.1 Implementation Details

We implement PPITrans with PyTorch based framework Fairseq (https://github.com/pytorch/fairseq). Embeddings from the ProtT5 (https://github.com/agemagician/ProtTrans) are of size 1024 and computed in advance.

We set the parameters of the PPITrans as follows while not tuning them elaborately. The Transformer encoder has 8 layers with hidden size being 256. There are 4 heads in the multi-head attention. PPITrans is trained 5 epochs with batch size 32 on two Nvidia Titan RTX GPUs. Its parameters are updated by Adam optimizer [44] with a learning rate of 3e-5. The dropout ratio is 0.2 in the training stage to avoid overfitting. Finally, we use the evaluation script provided by the scikit-learn (https://github.com/scikit-learn/scikit-learn) to compute the metrics in the paper. The threshold value of metrics, including Accuracy, Precision and Recall, is set to 0.5.

### 4.2 Comparison With Previous SOTA

We first compare PPITrans with recent state-of-the-art models on the cross-species binary PPI prediction task. There are four baselines, including PIPR [13], D-SCRIPT [10], PIPR+D-SCRIPT [10] (PIPR is used to adjust D-SCRIPT’s prediction), PIPR+S2F [29] (PIPR exploits the protein embeddings produced by S2F). All models are trained and tested on the dataset introduced in Section 3.2. We report metrics including precision, recall, F1 score, area under precision-recall curve (AUPR), and area under ROC curve (AUROC) of PPI prediction. Among these metrics, AUPR is generally considered a better metric for highly unbalanced data [10].

Table 3 lists the performance of all models on the PPIs of six species. It is evident that PPITrans outperforms other four models significantly, especially on the AUPR, which is generally considered a better metric for unbalanced data. In species other than Human, the improvement of PPITrans is more considerable. Even the most minor improvement of AUPR reaches 14.3 percentage points (on the PPIs of E.coli, from 0.588 to 0.731). Comparatively, the largest AUPR improvement reaches 33.9 percentage points (on the PPIs of Fly, from 0.562 to 0.901). A similar trend is also shown on the F1 score and AUROC. Such significant improvement accounts for the large recall ratio of PPITrans, as other models perform too poorly to identify positive samples (low recall ratio), although they have considerable precision. On the test set of Human PPIs, the performance gap of PPITrans over other models is relatively small. Despite all this, the AUPR of PPITrans still outperforms PIPR+D-SCRIPT, the best of previous models, by 8.3 percentage points. Throughout Table 3, PPITrans surpasses the four baselines evidently on all species and all metrics other than precision. However, the precision metric is less comprehensive in indicating the performance of models. Overall, PPITrans has a great advantage in generalization ability as it outperforms those baselines more significantly on PPIs of unseen species, indicating its superiority in alleviating the problem of OOD.

**TABLE 3.** Performance of models that are trained on human PPIs. Bold digits are the best on the subset.

| Species | Methods | AUPR | Precision | Recall | F1 | AUROC |
| --- | --- | --- | --- | --- | --- | --- |
| Mouse | PIPR | 0.526 | 0.734 | 0.331 | 0.456 | 0.839 |
|  | D-SCRIPT | 0.580 | 0.818 | 0.346 | 0.486 | 0.833 |
|  | PIPR+D-SCRIPT | 0.609 | **0.820** | 0.355 | 0.495 | 0.838 |
|  | PIPR+S2F | 0.644 | 0.781 | 0.447 | 0.568 | 0.889 |
|  | **PPITrans** | **0.895** | 0.785 | **0.844** | **0.813** | **0.978** |
| Fly | PIPR | 0.278 | 0.521 | 0.121 | 0.196 | 0.728 |
|  | D-SCRIPT | 0.552 | **0.798** | 0.359 | 0.495 | 0.824 |
|  | PIPR+D-SCRIPT | 0.562 | **0.798** | 0.361 | 0.497 | 0.824 |
|  | PIPR+S2F | 0.450 | 0.652 | 0.286 | 0.397 | 0.795 |
|  | **PPITrans** | **0.901** | 0.771 | **0.856** | **0.811** | **0.980** |
| Worm | PIPR | 0.346 | 0.673 | 0.142 | 0.235 | 0.757 |
|  | D-SCRIPT | 0.548 | 0.840 | 0.306 | 0.449 | 0.813 |
|  | PIPR+D-SCRIPT | 0.559 | 0.841 | 0.308 | 0.451 | 0.814 |
|  | PIPR+S2F | 0.484 | 0.724 | 0.270 | 0.393 | 0.825 |
|  | **PPITrans** | **0.870** | **0.863** | **0.740** | **0.797** | **0.969** |
| Yeast | PIPR | 0.230 | 0.398 | 0.085 | 0.140 | 0.718 |
|  | D-SCRIPT | 0.405 | 0.706 | 0.223 | 0.339 | 0.789 |
|  | PIPR+D-SCRIPT | 0.417 | **0.708** | 0.225 | 0.341 | 0.789 |
|  | PIPR+S2F | 0.356 | 0.604 | 0.180 | 0.393 | 0.767 |
|  | **PPITrans** | **0.718** | 0.681 | **0.639** | **0.660** | **0.916** |
| E.coli | PIPR | 0.308 | 0.629 | 0.131 | 0.217 | 0.675 |
|  | D-SCRIPT | 0.571 | 0.791 | 0.520 | 0.627 | 0.863 |
|  | PIPR+D-SCRIPT | 0.588 | **0.793** | 0.394 | 0.526 | 0.863 |
|  | PIPR+S2F | 0.371 | 0.467 | 0.309 | 0.372 | 0.748 |
|  | **PPITrans** | **0.731** | 0.630 | **0.665** | **0.647** | **0.917** |
| Human | PIPR | 0.835 | 0.838 | 0.701 | 0.763 | 0.960 |
|  | D-SCRIPT | 0.516 | 0.728 | 0.278 | 0.402 | 0.833 |
|  | PIPR+D-SCRIPT | 0.844 | **0.949** | 0.400 | 0.562 | 0.962 |
|  | PIPR+S2F | 0.822 | 0.836 | 0.672 | 0.745 | 0.960 |
|  | **PPITrans** | **0.927** | 0.851 | **0.861** | **0.856** | **0.984** |

By comparing AUPR, F1, or AUROC scores of PPITrans on five unseen species, we can find an interesting phenomenon: lower scores are associated with lower protein identites shown in Figure 1. For example, the Mouse and Fly scores are higher than those on the Yeast and E.coli. By inspecting the prediction results manually, we find that wrongly predicted proteins by PPITrans are typically more distant from the Human proteins. In this perspective, although PPITrans makes progress on unseen species, the OOD problem still needs further solutions.

To validate whether the performance of PPITrans is tightly related to the existence of homologous proteins to the Human training set, we re-assess the AUPR concerning the homology. Specifically, we gradually remove top homologous proteins in the test set of each species. As shown in Figure 4, the AUPR of PPITrans decreases with the identity decreasing and experiences a sharp degradation when the identity is smaller than 30%. However, only a few samples share an identity lower than 30%, except E.coli. It indicates that PPITrans performs poorly only on those samples of very low identities.

**Fig. 4.**
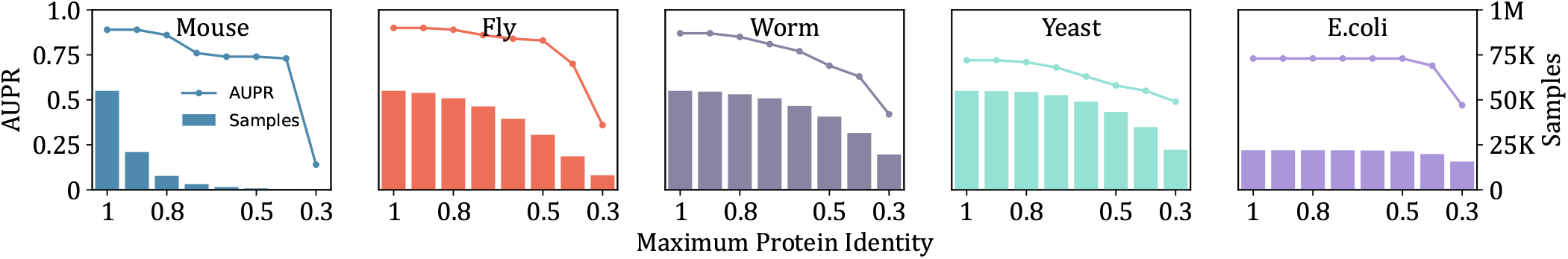
AUPR concerning protein identity. The histogram is the number of samples in which two protein identities with Human proteins are both smaller than the value on the horizontal axis. The line is the AUPR of PPITrans on the above samples.

### 4.3 Ablation Studies

More ablation studies are conducted to evaluate the contribution of each module to the significant performance of PPITrans. Specifically, we investigate two factors, the pretrained embedder and the Transformer encoder. We replace the pre-trained ProtT5 with a randomly initialized embedding layer and name the new model PPITrans-ProtT5. The embedding size of PPITrans-ProtT5 is controlled the same as ProtT5 for a fair comparison. To evaluate the contribution of the Transformer encoder, we remove the Transformer from PPITrans and name the new model as PPITrans-Transformer. Specifically, in PPITrans-Transformer, there is no protein encoder, so the classifier makes predictions based on the pre-trained protein embedding directly. Other settings of PPITrans-Transformer keep consistent with PPITrans.

Figure 5 shows the ablation study results. No surprise that PPITrans outperforms the other two models in all species and metrics, indicating that both pre-trained ProtT5 embedder and Transformer protein encoder are beneficial to PPI prediction. The gap between the three models is minimum on the Human PPIs, and PPITrans-ProtT5 exceeds PPITrans-Transformer on three metrics. However, PPI-Transformer overtakes PPI-ProtT5 largely on the PPIs of another five species. The result indicates that the prominent performance of PPITrans on unseen species accounts largely for ProtT5 rather than Transformer, as the training data of ProtT5 could cover all proteins. Therefore, it is possible that PPITrans is able to generalize to more unseen species. On the other hand, Transformer also plays an essential role in learning interactions between two proteins because PPITrans outperforms PPITrans-Transformer on all species, especially the Human.

**Fig. 5.**
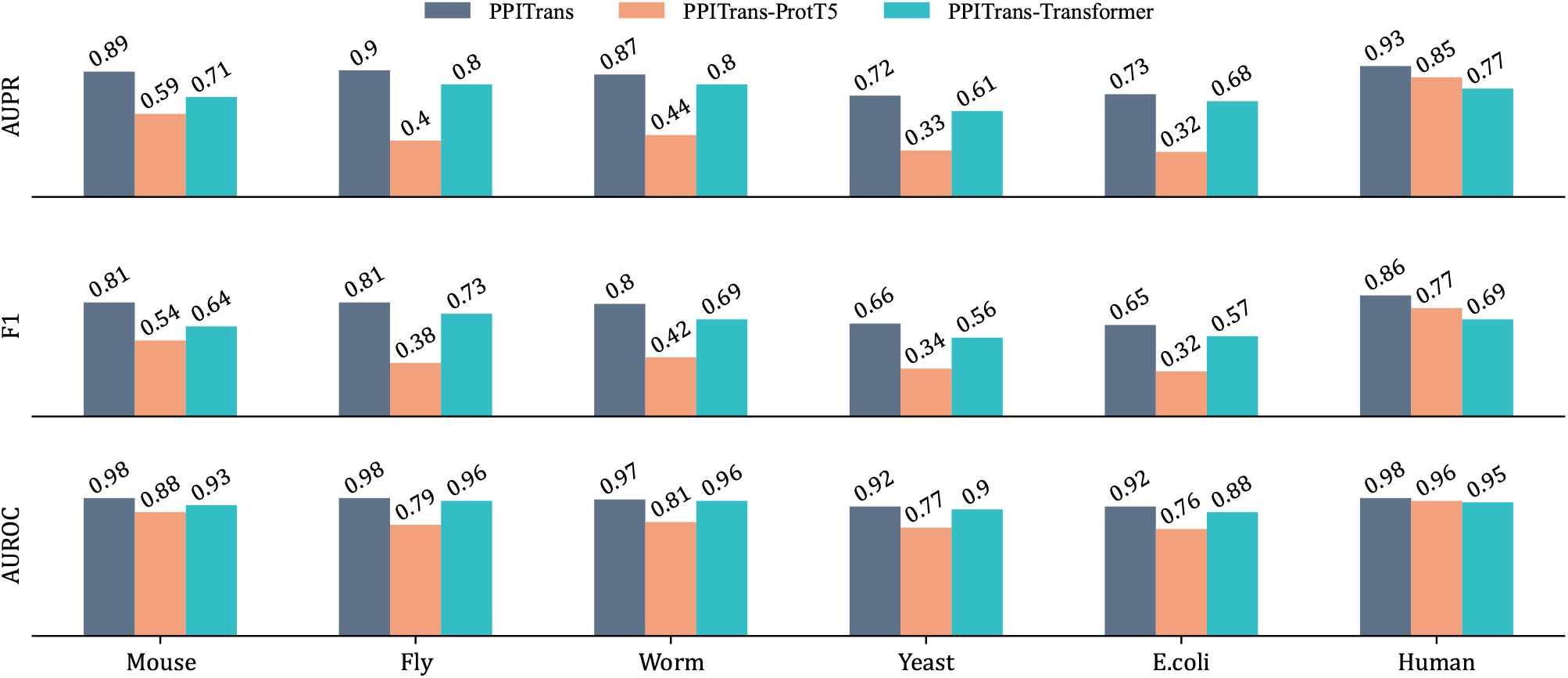
Results of ablation studies.

To verify the separability of protein pair representations, we take protein representations output by each layer of PPITrans for PPI prediction. Specifically, for PPIs of specific species, we exploit ten KNNs (nearest five neighbors) to classify the Hadamard product of two protein representations (Equation 6) produced by all layers.s We randomly choose 5,000 PPIs from the test set of six species, of which 4,000 are for training while the rest 1,000 are splited into 5 folds for testing. Figure 6 plots the F1 scores of all KNNs for each layer and species. The output of the embedder layer is the most inseparable in five of six species, indicating that it is necessary to utilize the Transformer encoder. For species like Mouse and Human, the protein representations become more and more separable from the bottom to the top layer. However, deep encoder does not always bring better separability. For example, the separability of top layers keeps flat or even decreases slightly for species like Fly, Worm and E.coli. The potential reason lies that the PPIs of these species are more distant from the training data of PPITrans (Human PPIs).

**Fig. 6.**
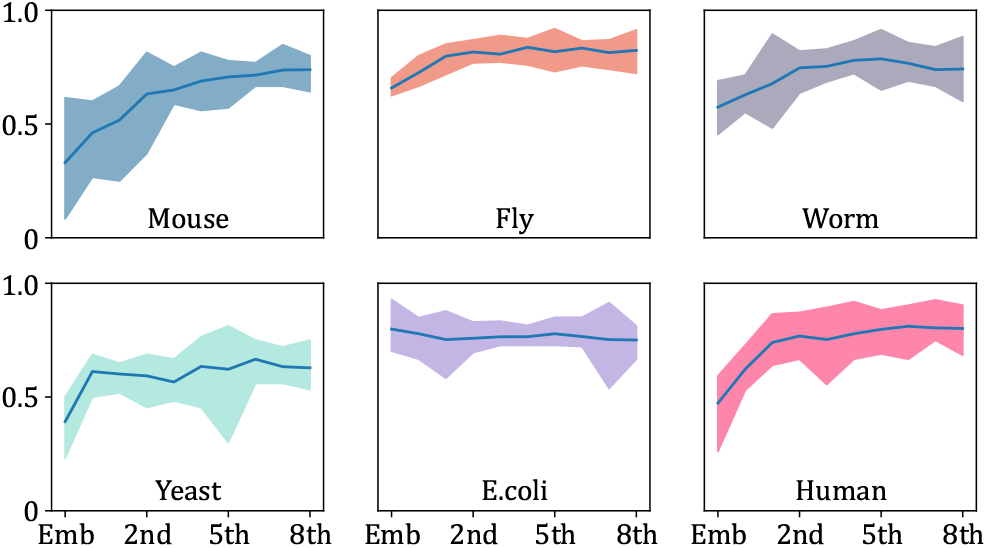
Performance of KNN classifiers for PPI prediction. The horizontal axis represents ten layers of PPITrans, and the vertical axis represents F1 scores. Each subplot reports the results of ten classifiers in specific species. Color blocks indicate the minimum and maximum values, and the line indicates the average.

### 4.4 Corrupt Training Set

We corrupt the Human training set and test the PPITrans on the original test sets. Specifically, the training set is corrupted in two manners: reducing the training set, ranging from 80% to 20%, and replacing positive samples with randomly selected protein pairs, ranging from 20% to 100%. Figure 7 shows the performance of PPITrans trained on the corrupted Human training set.

**Fig. 7.**
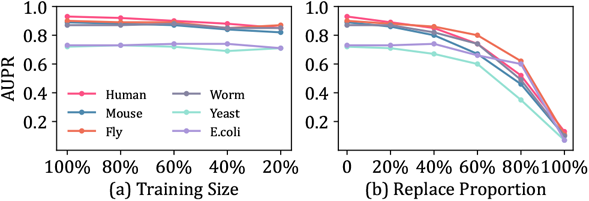
The performance of PPITrans trained on corrupted training sets. (a) The training set is reduced; (b) Proportional positive samples in the training set are replaced with random protein pairs.

As shown in Figure 7(a), the size of the training set has little impact on the model performance. Even though only 20% samples are reserved, the PPITrans still obtains competent AUPR as trained on the complete training set, surpassing previous models vastly. It implies that the outstanding performance PPITrans does not result from the large training set size. The result in Figure 7(b) shows a surprising phenomenon that PPITrans would perform well unless all positive samples are replaced. For example, when replacing 80% positive samples, PPITrans can still outperform PIPR. In conclusion, Figure 7 proves that PPITrans is robust to learning from small datasets, even those of low quality.

### 4.5 Results of Residue Level Probes

Figure 8 shows the accuracy of ten KNNs in the probe of amino acid classification. All curves in six species display an almost identical trend: from the bottom to the top layers, the identifiability of amino acids decreases. The embedder layer’s high accuracy lies in that ProtT5 adopts amino acid prediction as one of its pre-training tasks. Besides, the variance of the accuracy of five folds (width of lines in Figure 8) is pretty tiny, indicating that the classifier is stable to make predictions. As for the second probe at the residue level, Figure 9 shows the same phenomenon as in Figure 8: it becomes more difficult to predict the physicochemical properties from the bottom to the top. The results of these two probes at the amino acid level indicate that features of both amino acids and their physicochemical property vanish gradually. Furthermore, such a phenomenon displays high consistency across all species. They prove that PPITrans is incapable of retraining fine-grained amino acid level features. But PPITrans also gradually extracts coarse-grained features for PPI prediction.

**Fig. 8.**
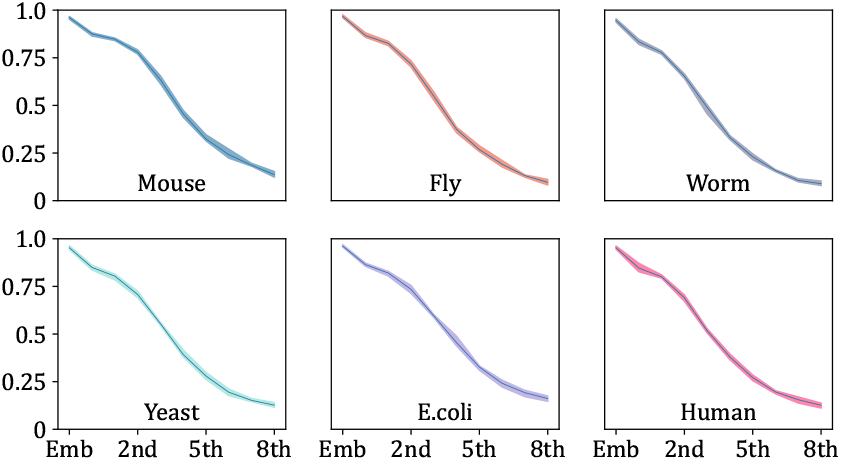
Accuracy of ten KNN classifiers in the probe of amino acid classification.

**Fig. 9.**
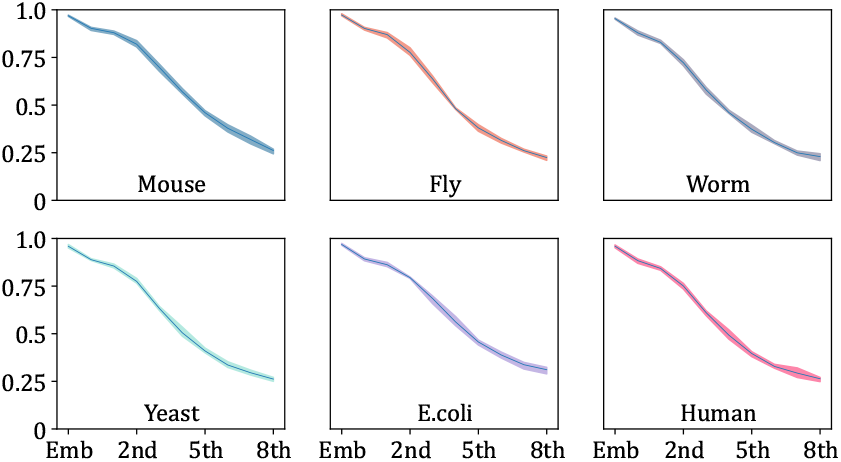
Accuracy of KNN classifiers in the probe of physicochemical property classification.

### 4.6 Results of Protein Level Probes

Table 4 reports accuracies and standard deviations of the probe of secondary structure prediction at the protein level. At the bottom layers, the secondary structure prediction accuracy is relatively appreciable, especially at the projection layer that reaches 0.75. It shows that the amino representations output by these layers can be easily separated in the perspective of the secondary structure. However, the accuracy gradually decreased to 0.421, 0.329 less than the maximum value. This evidence indicates that PPITrans cannot retain secondary structure information. In other words, the secondary structure is not the crucial factor leveraged by the PPITrans, or even not required by the PPITrans for PPI prediction.

**TABLE 4.** Accuracy of the secondary structure prediction probe.

| Layer | Accuracy | Layer | Accuracy |
| --- | --- | --- | --- |
| Emb | 0.706±0.020 | 4th | 0.512±0.018 |
| Proj | 0.750±0.020 | 5th | 0.477±0.018 |
| 1st | 0.721±0.016 | 6th | 0.440±0.010 |
| 2ec | 0.658±0.020 | 7th | 0.433±0.009 |
| 3rd | 0.590±0.029 | 8th | 0.421±0.009 |

Since PPITrans fails to grasp the secondary structure, a natural question is that will it capture the spatial structure of proteins, which can be represented as a contact map. So we devise the second probe for the contact map prediction. As the unbalance of data, we report the F1 scores of ten KNN classifiers in Table 5. Unfortunately, the probe of contact map prediction displays a similar result to the one for the secondary structure. For example, the maximum F1 score lies in the 1st layer but decreases from 0.361 to 0.162. Therefore, PPITrans is incapable of capturing the spatial structure of proteins.

**TABLE 5.** F1 scores of the contact map prediction probe.

| Layer | F1 | Layer | F1 |
| --- | --- | --- | --- |
| Emb | 0.351±0.016 | 4th | 0.241±0.016 |
| Proj | 0.318±0.011 | 5th | 0.206±0.014 |
| 1st | 0.361±0.008 | 6th | 0.203±0.006 |
| 2ec | 0.352±0.012 | 7th | 0.198±0.009 |
| 3rd | 0.300±0.027 | 8th | 0.162±0.006 |

Figure 10 shows the performance of all ten KNNs in the probe of species classification. It displayers two interesting findings. Firstly, F1 scores of six species have an obvious hierarchy, indicating that proteins of some species are easier to identify, such as E.coli and Yeast. These two species are also more genetically distant from the Human. Secondly, although F1 scores of all species experience a slight decrease from the bottom to the top, the drop in E.coli and Yeast is more evident. These results imply that protein representations of all species tend to be homogenous from the bottom to the top. On the other hand, the result may also prove that the ability of the model to alleviate the OOD problem gradually improves from the bottom layer to the top layer.

**Fig. 10.**
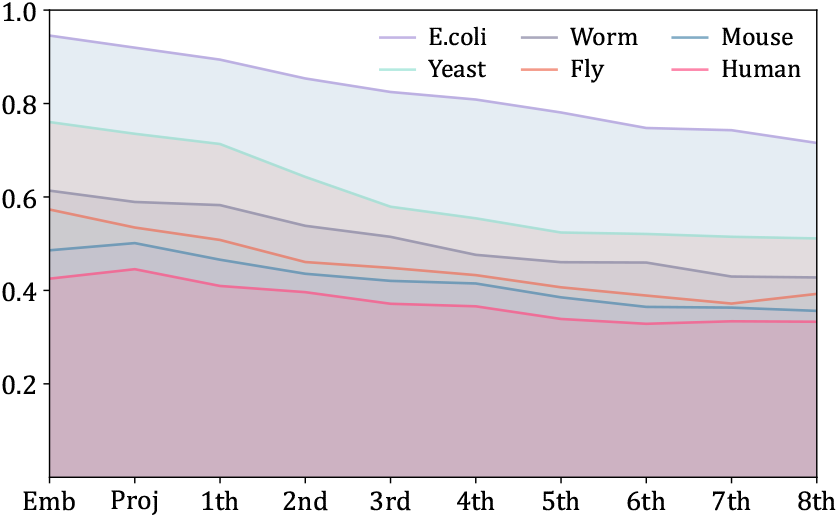
F1 scores of ten KNN classifiers for species classification probe.

### 4.7 Results of Protein Pair Level Probes

In the probe of contact map prediction at the protein pair level, the KNN classifier takes the Hadamard product of two residue representations and predicts whether they contact or not. F1 scores of all ten KNNs for protein pair contact map prediction are shown in Table 6. Like results in Table 5, the F1 score reaches the maximum at the 1st layer and decreases until the last layer, although the decline shown in Table 6 is minor. Moreover, as shown in Figure 11, we conduct a visualization of the gold contact map and top residue pairs that contribute the most to the interaction likelihood (Equation 9). Figure 11 shows that many residue pairs that contribute the most to likelihood are not in contact. It seems that PPITrans is inclined to focus on the relationship between some particular residues pairs, rather not those in contact.

**Fig. 11.**
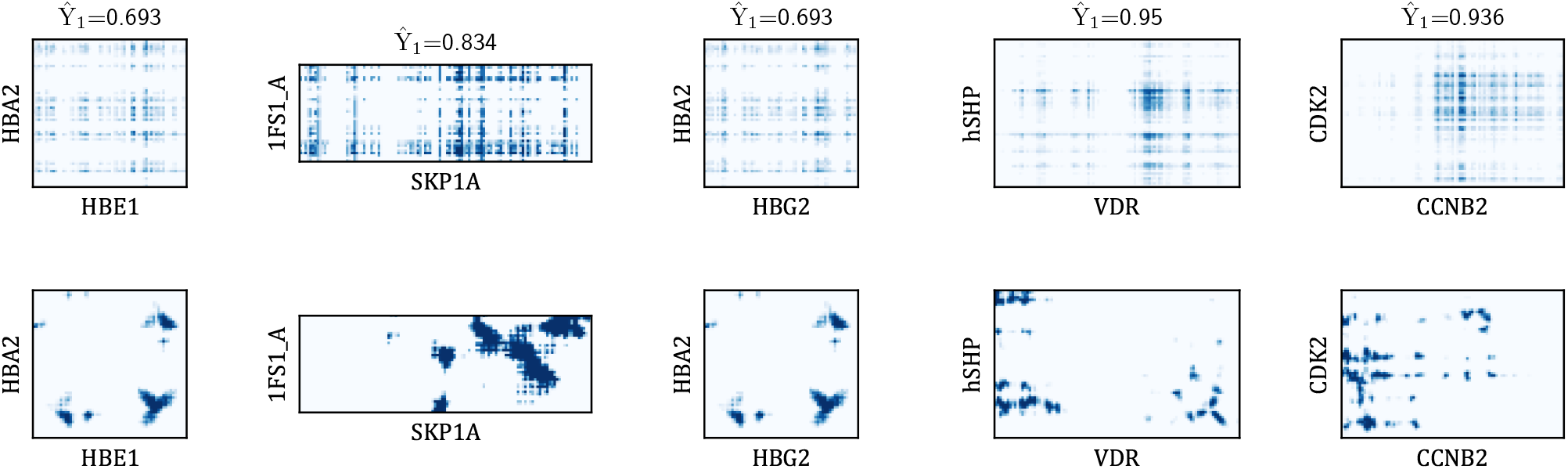
Cases of protein pair contact map. The lower row are labled samples from GWIDD, while the upper row are visualization of top residue pairs contributing the most to the interaction likelihood 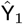. We control the number of contacted residue pairs at the upper the same as the lower.

**TABLE 6.** F1 scores of protein pair contact map prediction probe.

| Layer | F1 | Layer | F1 |
| --- | --- | --- | --- |
| Emb | 0.394±0.023 | 4th | 0.432±0.014 |
| Proj | 0.445±0.013 | 5th | 0.420±0.018 |
| 1st | 0.457±0.007 | 6th | 0.394±0.020 |
| 2ec | 0.454±0.006 | 7th | 0.366±0.013 |
| 3rd | 0.451±0.014 | 8th | 0.329±0.013 |

The second probe at the protein pair level is to verify whether PPI type can be predicted from protein pair representations. Table 7 shows the performance of ten KNNs for PPI type prediction. In general, the divergence in the performance of KNNs in different layers is slight. However, the accuracy of KNNs in the last layer outperforms the one in the embedder layer with about 3 percentage points, indicating PPITrans has learned knowledge about PPI types.

**TABLE 7.** Accuracy of PPI type prediction probe.

| Layer | Accuracy | Layer | Accuracy |
| --- | --- | --- | --- |
| Emb | 0.406±0.016 | 4th | 0.433±0.005 |
| Proj | 0.410±0.005 | 5th | 0.422±0.014 |
| 1st | 0.421±0.013 | 6th | 0.423±0.008 |
| 2ec | 0.427±0.016 | 7th | 0.423±0.004 |
| 3rd | 0.421±0.013 | 8th | 0.431±0.018 |

The third probe at the protein pair level is to estimate the binding affinity of interacted protein pairs. Table 8 shows the MSE loss of ten estimators on the test set. We are surprised to find that the MSE loss degrades sharply from 8.134×10^-2^ to 2.939×10^-2^. It indicates that the pretrained embedding produced by ProtT5 performs worse on estimating binding affinity, while PPITrans is capable of obtaining such ability, although it is trained only for binary PPI prediction.

**TABLE 8.** Mean square loss of binding affinity estimation probe.

| Layer | MSE ($\times 10^{-2}$) | Layer | ($\times 10^{-2}$) |
| --- | --- | --- | --- |
| Emb | 8.134±1.000 | 4th | 2.943±0.682 |
| Proj | 3.618±0.797 | 5th | 2.843±0.509 |
| 1st | 3.303±0.447 | 6th | 2.862±0.587 |
| 2ec | 3.020±0.601 | 7th | 2.917±0.406 |
| 3rd | 3.044±0.342 | 8th | 2.939±0.452 |

The experimental results at the protein pair level firstly indicate again that PPITrans does not make predictions through the spatial structure of proteins. However, another two probes at the protein pair level also reveal an inspiring conclusion that PPITrans is able to go beyond binary PPI prediction and learn some knowledge about PPI type and binding affinity.

## 5 Conclusion

In this paper, we propose PPITrans for binary PPI prediction. To improve prediction performance on unseen species, PPITrans exploits a language model pre-trained on large-scale protein datasets to promote PPI prediction across species. Experimental results show that PPITrans outperforms the previous state-of-the-art on the Human PPIs and generalize better significantly on other species. Especially on the Fly PPIs, the AUPR improvement reaches surprising 33.9 percentage points. The superior performance of PPITrans stimulates us to explore what PPITrans has learned. Therefore, we devise various probes at three levels to verify the knowledge learned by PPITrans. At the amino acid level, two probes reveal PPITrans extracts globally coarse-grained features while dropping fine-grained amino acid features. The three probes at the protein level indicate PPITrans does not rely on protein structure to make predictions, although it performs well across species. Finally, the first probe at the protein pair level proves again that the superiority of PPITrans does not originate from identifying protein structures. However, other probes reveal that PPITrans go beyond binary PPI prediction and learn the knowledge of PPI type and binding affinity automatically. In view of the results of these probes, a future research direction may be driving the PPI prediction model learns protein structure automatically so as to shed more light on its interpretability.

## Availability of Data and Codes

All datasets and source codes in this study are available at https://github.com/LtECoD/PPITrans.

## Acknowledgments

This work is supported in part by funds from the National Natural Science Foundation of China (81830101).

